# Fast and specific enrichment of cancer-related exosomes by DNA-nanoweight-assisted centrifugation

**DOI:** 10.1101/2021.12.28.474334

**Authors:** Zhangwei Lu, Ye shi, Yuxuan Ma, Bin Jia, Xintong Li, Zhe Li

## Abstract

Exosomes are nanoscale membrane vesicles actively released by cells and play an important role in the diagnosis of cancer-related diseases. However, it is challenging to efficiently enrich exosomes from extracellular fluids. In this work, we used DNA-tetrahedron as a nanoweight during centrifugation to precisely enrich tumor exosomes from a complex biological environment. Two different DNA tetrahedral nanostructures (DTAs), each carrying a specific aptamer for exosome biomarker recognition, were incubated with clinical samples simultaneously. One DTA triggered the cross-linking of multiple target exosomes, and therefore enabled low-speed and fast centrifugation for enrichment. The other DTA further narrowed down the target exosome subtype and initiated a hybridization chain reaction (HCR) for sensitive signal amplification. The method enabled the detection of 180 MCF-7-derived exosomes per microliter and 560 HepG2-derived exosomes per microliter, with 1000-fold higher sensitivity than conventional ELISA. This easy-to-operate method can enrich exosomes with excellent specificity and therefore will be appealing in biomedical research and clinical diagnosis.

## Introduction

Exosomes, nanosized extracellular vesicles actively released from cells, play an indispensable role in physiological activities^1,2^. In particular, exosomes derived from cancer cells are involved in mediating intercellular communication to facilitate cancer growth and metastasis^3–5^. It has been reported that the expression of exosomal membrane proteins such as CD8, CD63, EpCAM, PCAT-1 and PSA is much higher on cancerous exosomes than on noncancerous exosomes^5,6^. Therefore, exosomes have been proposed as biomarkers for the early noninvasive diagnosis of cancer^7,8^.

Although exosomes are potentially valuable for clinical diagnosis, their small size and complex biological environment hinder their rapid separation. The current gold standard of exosome separation is ultracentrifugation^9^. However, the bulky and costly instrumentation, time-consuming process, and requirement of a large number of samples limit its applications. Other separation methods have been reported, including size exclusion chromatography^10,11^, microfluidic filtration^12,13^ and polymer-based precipitation^14,15^, but they suffer from low purity, irreversible membrane destruction, or complex operation. Therefore, it is still urgent to establish convenient methods to enrich tumor-derived exosomes in routine clinical testing.

Herein, we used DNA-tetrahedron as a nanoweight during centrifugation to achieve this goal. We designed two DNA tetrahedral nanostructures (DTAs), each carrying a specific aptamer sequence. One DTA served as the nanoweight for exosomes, with one vertex modified with an aptamer to recognize the typical exosomal protein CD63^22^ and the other three with capture strands to cross-link multiple target exosomes. The large nanoparticles formed from cross-linked exosomes and the high buoyant density of DTA allowed enrichment under a low centrifugal speed (20,000×g) with a short time period (30 min). The other DTA acted as the signal amplifier, with one vertex modified with another aptamer targeting a tumor-specific marker (AFP^23^ or EpCAM^24^) and the other three with toeholds to trigger a hybridization chain reaction (HCR). The use of two aptamers performed a more accurate survey of the exosomal surface, and the HCR provided in situ signal amplification, significantly reducing the interfering signals. Our method enabled the detection of 1.8×10^2^ MCF-7-derived exosomes per microliter and 5.6×10^2^ HepG2-derived exosomes per microliter.

## Results and Discussion

### System design

Inspired by a recent study that used DNA nanostructures to assist the separation of liposomes of different sizes^25^, we designed a DNA tetrahedron as a nanoweight (Weight-DTA, Figure 1) to facilitate the enrichment of exosomes and another DNA tetrahedron as a HCR amplifier (HCR-DTA). The DTA nanostructures were designed from 4 ssDNA sequences as previously reported^26^.

**Figure 1.**
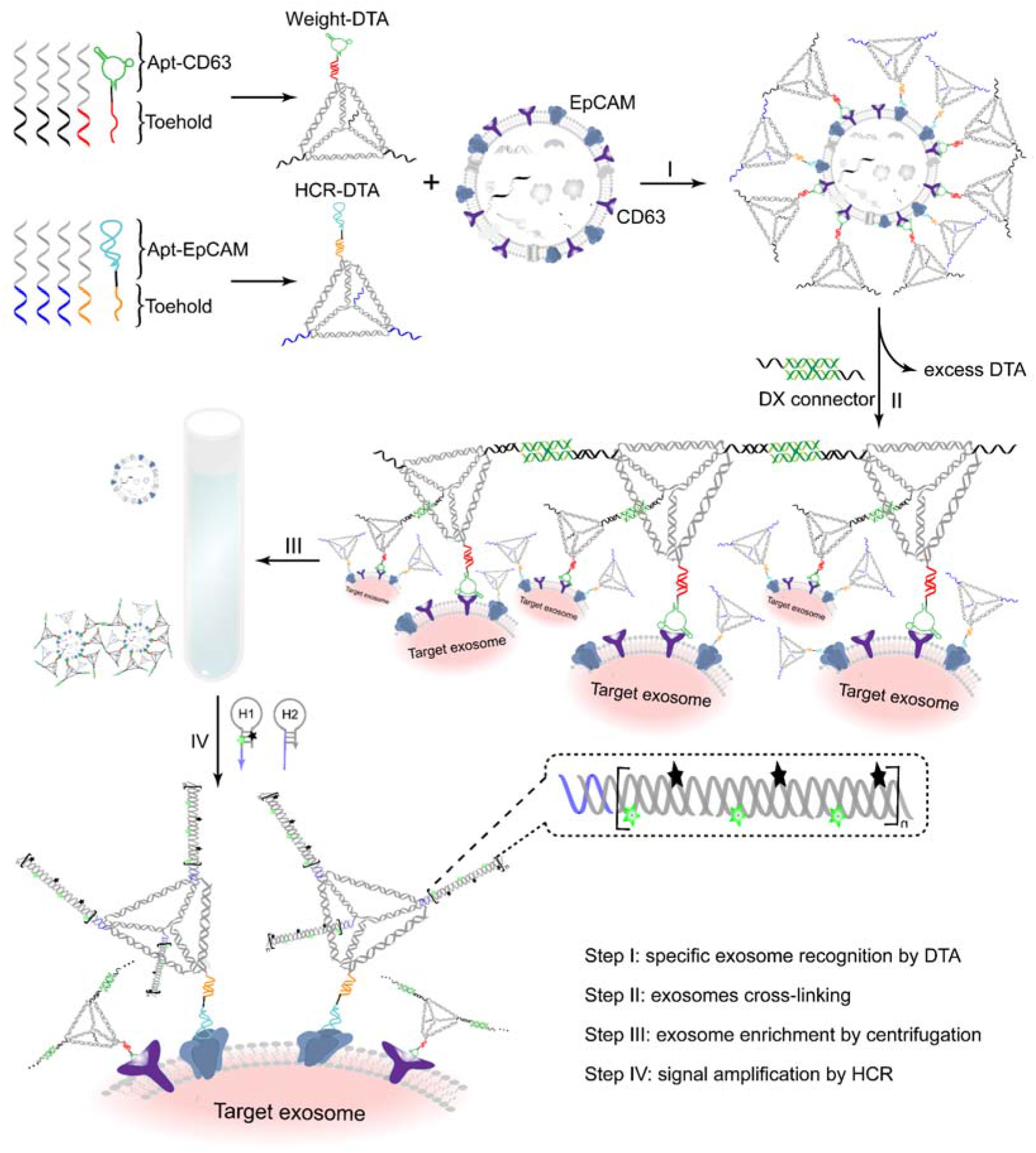
Schematic illustration of DNA-tetrahedron-assisted centrifugation for exosome enrichment. (A) Structure of the DNA tetrahedron as the nanoweight. (B) Structure of the DNA tetrahedron as the HCR amplifier. (C) Steps of exosome enrichment.

The Weight-DTA comprises two active components. The first is the CD63 aptamer (Apt-CD63), which is hybridized to DTA through an extended strand from one vertex. The second is an ssDNA linker that extends from each of the other vertexes. The linker can cross-link with each other in the presence of the DX connector, a two double-helix structure formed by two partially complementary ssDNAs. Similarly, HCR-DTA also contains a specific aptamer at one vertex. It is either the EpCAM aptamer (Apt-EpCAM) or AFP aptamer (Apt-AFP) to specifically recognize MCF-7 or HepG2 cell-secreted exosomes. Each of the other three vertexes contains an initiator sequence of HCR.

The specific recognition of aptamers leads to the anchor of Weight-DTA and HCR-DTA. The use of two aptamers in this design narrows down the subpopulations of exosomes with different expression patterns of exosomal membrane markers, enabling accurate identification of exosomes. Next, with the help of a DX connector, multiple exosomes are cross-linked, which allows their separation by low-speed centrifugation. After centrifugation at 20,000×g for 30 minutes, the crosslinked exosomes are isolated as pellets. Then two probes, FAM-H1-BHQ and H2, are added, leading to the trigger of the HCR. An amplified fluorescence signal is activated in the chain-like HCR product due to the separation of fluorophore FAM from quencher BHQ^27^.

### Characterization of exosome enrichment

The assembly of Weight-DTA (Figure 2A) and HCR-DTA (Figure 2B) was first evaluated by 6% native PAGE. The assembled structures showed clear master bands on the gel, and the migration rate of final products was slower than that of each component strand. HCR were also evaluated by native PAGE (Figure 2C). When only F-H1-Q and H2 were mixed, no new band could be observed, indicating that the two strands coexisted stably. Upon the addition of HCR-DTA, multiple large products appeared, suggesting the successful triggering of HCR.

**Figure 2.**
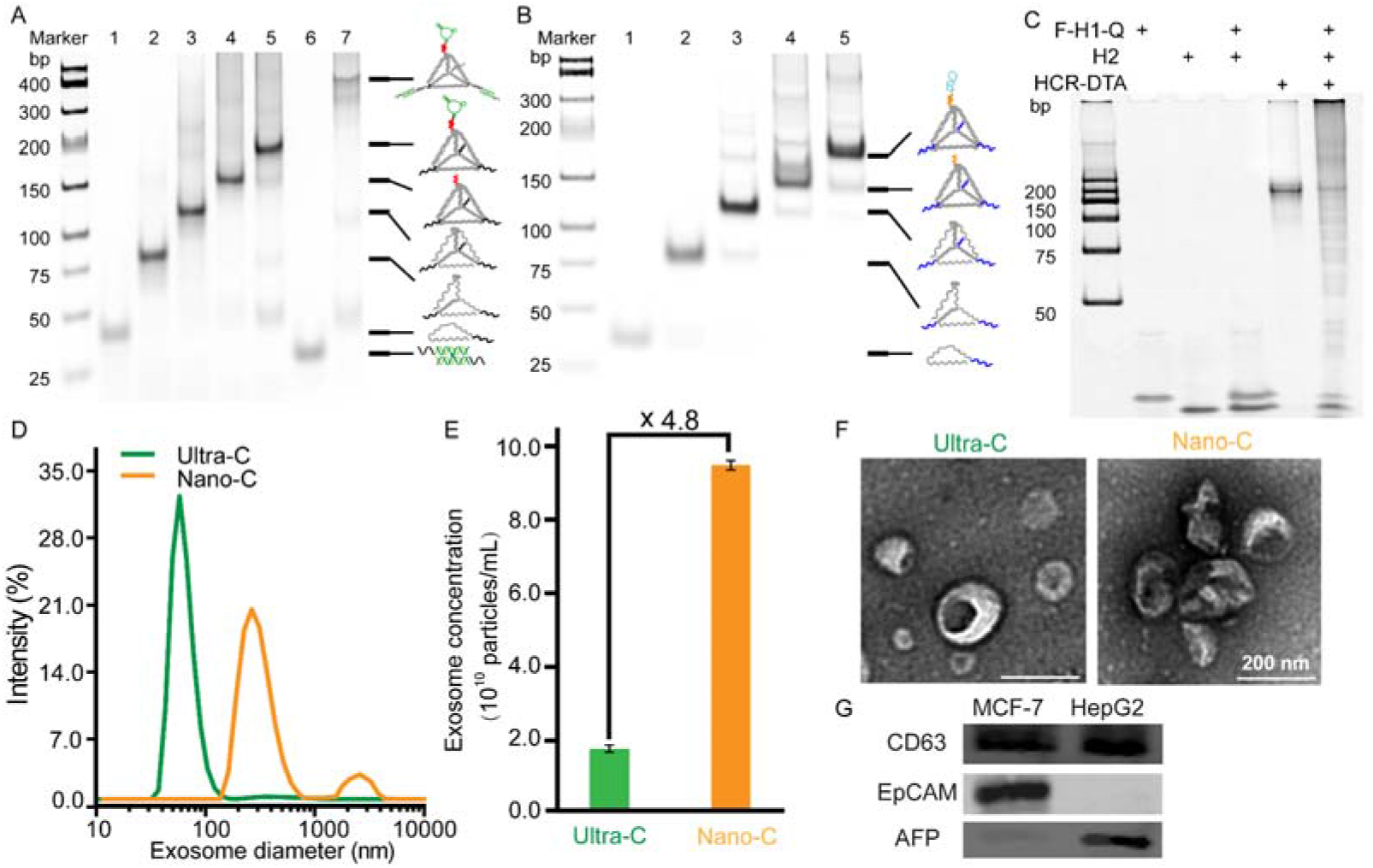
Enrichment of MCF-7-derived exosomes by Nanoweight-assisted centrifugation (Weight-C). (A) 6% native PAGE analysis of the gradual formation of Weight-DTA. Lane 1-4: the mixture of one, two, three, and four components strands of DTA respectively. Lane 5: DTA + Apt-CD63. Lane 6: DX connector. Lane 7: DTA + DX connector. (B) 6% native PAGE analysis of the gradual formation of HCR-DTA. Lane 1-4: the mixture of one, two, three, and four components strands of DTA respectively. Lane 5: DTA + Apt-EpCAM. (C) 6% native PAGE analysis of the gradual formation of HCR. (D) DLS analysis, (E) total protein amount, (F) concentration, and (G) TEM images of MCF-7-derived exosomes isolated by Weight-C or conventional ultracentrifugation (U-C). Data are represented as the means ± SD (n = 4). (H) Western blotting analysis of CD63, AFP and EpCAM proteins of MCF-7- and HepG2-derived exosomes enriched by COAT-DTA.

Crude exosomes were first isolated from MCF-7 cells by centrifugation and then incubated with Weight-DTA and HCR-DTA/Apt-EpCAM. The addition of the DX connector caused cross-linking of multiple exosome particles and allowed isolation through low-speed centrifugation (20,000×g for 30 min). As a comparison, conventional ultracentrifugation (U-C) was performed twice at 140,000×g for 90 min to enrich the crude sample.

The exosomes obtained from both methods were measured by dynamic light scattering (DLS). The particle size increased significantly from 80 nm to 300 nm after the addition of DTA and the DX connector, and a small aggregate peak appeared above 1 μm, indicating the cross-linking of multiple exosome particles (Figure 2D). The total protein content and concentration of exosomes enriched from both methods were characterized through bicinchoninic acid (BCA) quantification and nanoparticle tracking analysis (NTA). Compared with conventional ultracentrifugation, DTA-assisted centrifugation increased the total protein content by 2.89-fold (Figure 2E) and the concentration by 4.8-fold (Figure 2F). Exosome particles were then directly visualized by TEM. While those from ultracentrifugation seemed monodisperse, those from DTA-assisted centrifugation appeared aggregated with good structural integrity (Figure 2G).

To validate the generality of the method, we then incubated Weight-DTA and HCR-DTA/Apt-AFP with crude HepG2-derived exosomes. The particle size, total protein content, concentration, and morphology of HepG2 exosomes were analyzed and showed similar results (Figure S1). All the above results indicated that DNA nanostructures performed impressively for exosome enrichment. The expression of CD63, EpCAM and AFP by the enriched exosomes was verified using Western blotting analysis. Clear bands of CD63 were observed in both samples, whereas EpCAM and AFP were only observed in MCF-7- or HepG2-derived exosomes, respectively (Figure 2H), corroborating previous evidence on exosome characterization^28–30^.

### Characterization of exosome quantification

After DNA-nanoweight-assisted centrifugation, excess DTAs were removed, and F-H1-Q and H2 were added to trigger the HCR. For MCF-7-derived exosomes, the resulting fluorescence intensity significantly increased by 3.6-fold compared to that without HCR (Figure 3A). Meanwhile, exosomes enriched from ultracentrifugation were treated in the same way, and a 2.5-fold increase in fluorescence intensity was obtained (Figure 3B), indicating that HCR improved the detection sensitivity. Interestingly, exosomes enriched by ultracentrifugation and Weight-DTA-assisted centrifugation displayed much different fluorescence intensities after triggering HCR. The enrichment of Weight-DTA greatly increased the signal by 6.8-fold (Figure 3C). Similarly, For HepG2-derived exosomes, a 3.2-fold (U-C) or 2.4-fold (Weight-C) higher fluorescence intensity was detected. In addition, compared with conventional ultracentrifugation, Weight-DTA-assisted centrifugation increased the signal by 5.4-fold (Figure S2). The results indicated that the introduction of both DNA nanoweight and HCR amplification greatly improved the detection sensitivity.

**Figure 3.**
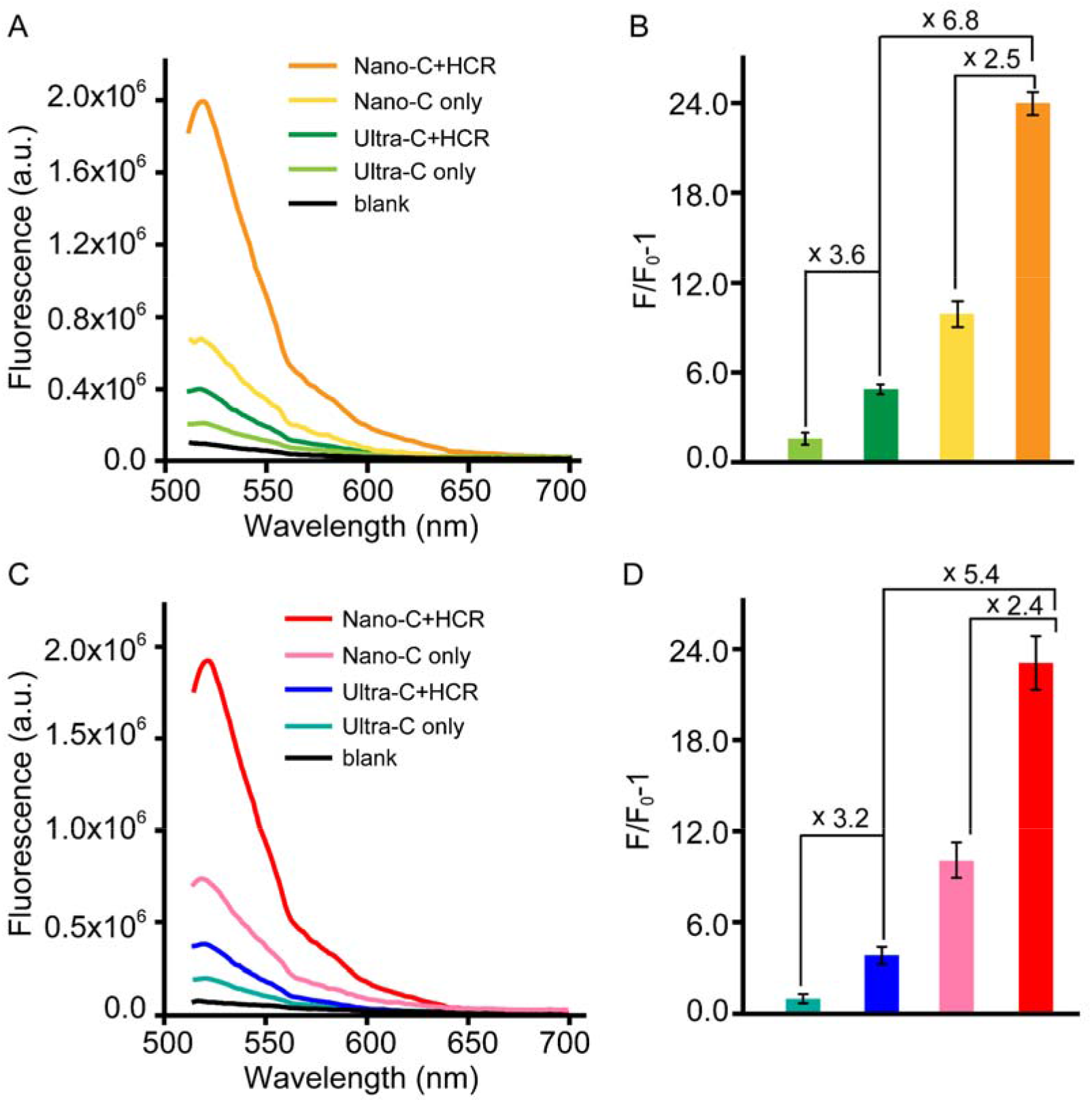
Quantification of MCF-7-derived and HePG2-derived exosomes by HCR. (A) Fluorescence spectrum of MCF-7-derived exosomes enriched from Ultra-C or Nano-C with or without (w/o) HCR. (B) Fluorescence enhancement ratios (F/F_0_ - 1) corresponding to the data from (A). (C) Fluorescence spectrum of MCF-7-derived exosomes enriched from Ultra-C or Nano-C with or without (w/o) HCR. (D) Fluorescence enhancement ratios (F/F_0_ - 1) corresponding to the data from (C). Data are represented as the means ± SD (n = 3) (*** p < 0.005).

The limit of detection was then investigated. First, after enrichment by Weight-DTA and HCR-DTA/Apt-EpCAM, different concentrations of MCF-7-derived exosomes (measured by NTA) were incubated with H1 and H2, and the fluorescent signals after HCR were recorded (Figure 4A). A linear relationship was obtained between the fluorescent change ratio and logarithm of exosome concentrations, from concentrations of 1.7×10^4^ to 1.7×10^10^ particles/μl (Figure 4B). The limit of detection (LOD) was calculated to be 1.8×10^2^ particles/μl. Meanwhile, the same HCR-DTA/Apt-EpCAM was incubated with HepG2 exosomes, but the corresponding fluorescent signal was 6.9-fold lower (Figure 4C), demonstrating the high specificity of the method.

**Figure 4.**
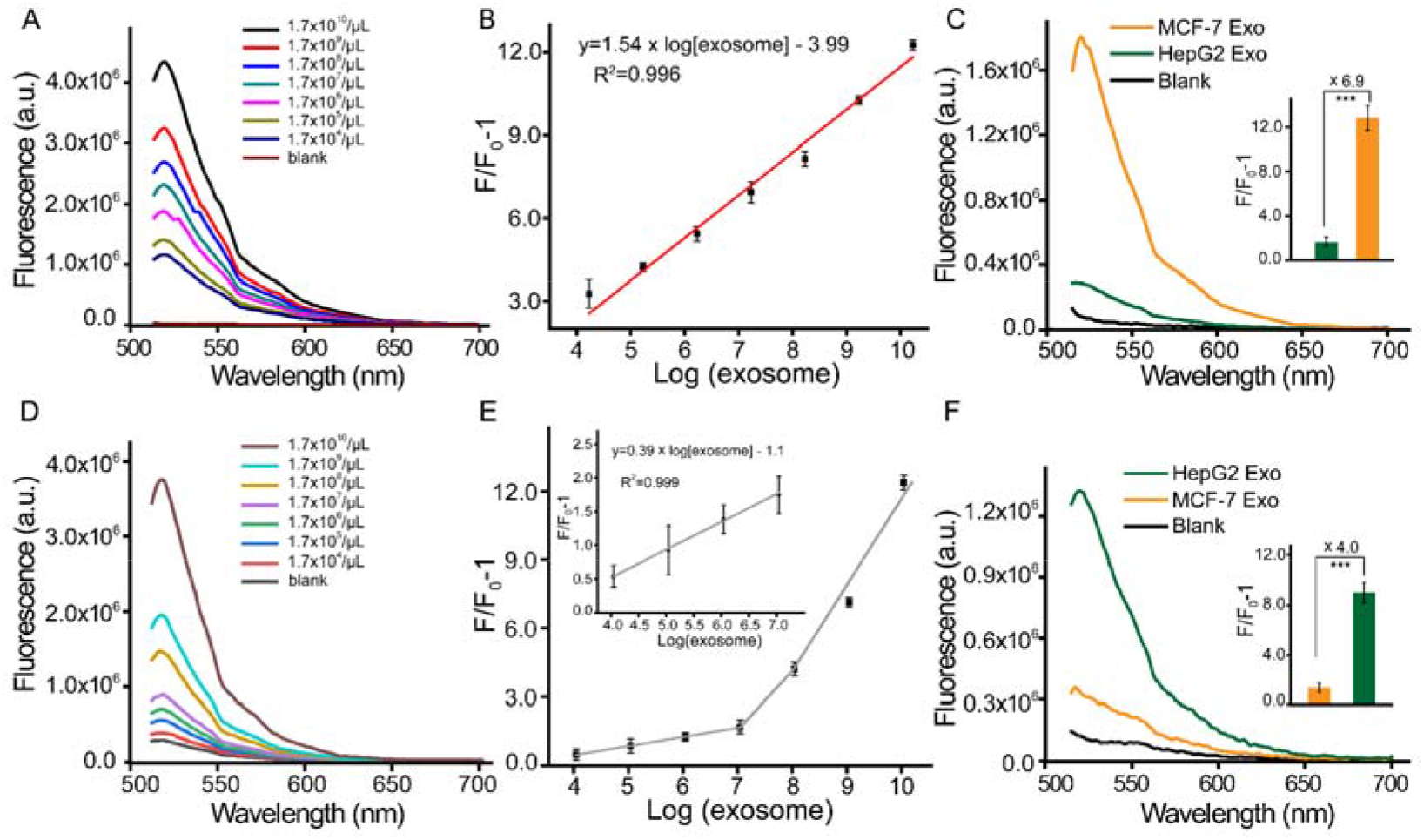
Sensitivity and specificity investigation of the proposed assay for exosome detection. (A) Fluorescence spectrum obtained at different concentrations of MCF-7 exosomes. (B) Linear plot of fluorescence intensity as a function of the base-10 logarithm (log for short) of the MCF-7 exosome concentration from 1.7×10^4^ to 1.7×10^10^ particles/μL. (C) Fluorescence spectrum of the detection of exosomes derived from MCF-7 and HepG2 cells by the same HCR-DTA. (D) Fluorescence spectrum obtained at different concentrations of HepG2 exosomes. (E) Linear plot of fluorescent intensity as a function of the base-10 logarithm of the HepG2 exosome concentration from 1.1×10^4^ to 1.1×10^7^ particles/μL. (F) Fluorescence spectrum of the detection of exosomes derived from MCF-7 and HepG2 by the same HCR-DTA. The error bars represent the standard deviations of three repetitive measurements (*** p < 0.005).

Similarly, HepG2-derived exosomes were enriched by Weight-DTA and HCR-DTA /Apt-AFP, and HCR was triggered by H1 and H2. A linear relationship was also generated between the fluorescent signal and the logarithm of HepG2 exosome concentrations, in the range of 1.1×10^4^ to 1.1×10^7^ particles/μL with a high correlation efficiency (Figure S3). The LOD was calculated to be 5.6×10^2^ particles/μl. To investigate the specificity, the same AMPL-DTA/Apt-AFP was incubated with MCF-7 exosomes; however, the resulting fluorescence intensity was 4.02-fold lower. The LOD of our method was comparable with several previously reported methods (Table S1). At the same time, our method featured high sensitivity, high specificity and simple operations.

### Clinical Application

Finally, our method was employed to analyze clinical samples. Nine freshly collected serum samples were tested, two from healthy individuals (H1 and H2) and the others from breast cancer patients (P1-P7). The concentration of MCF-7-derived exosomes from cancer patients was approximately twofold higher than that from normal controls (Figure 5A and 5B), demonstrating that our proposed method was suitable for the detection of clinical serum samples and could effectively distinguish breast cancer patients from healthy control testers.

**Figure 5.**
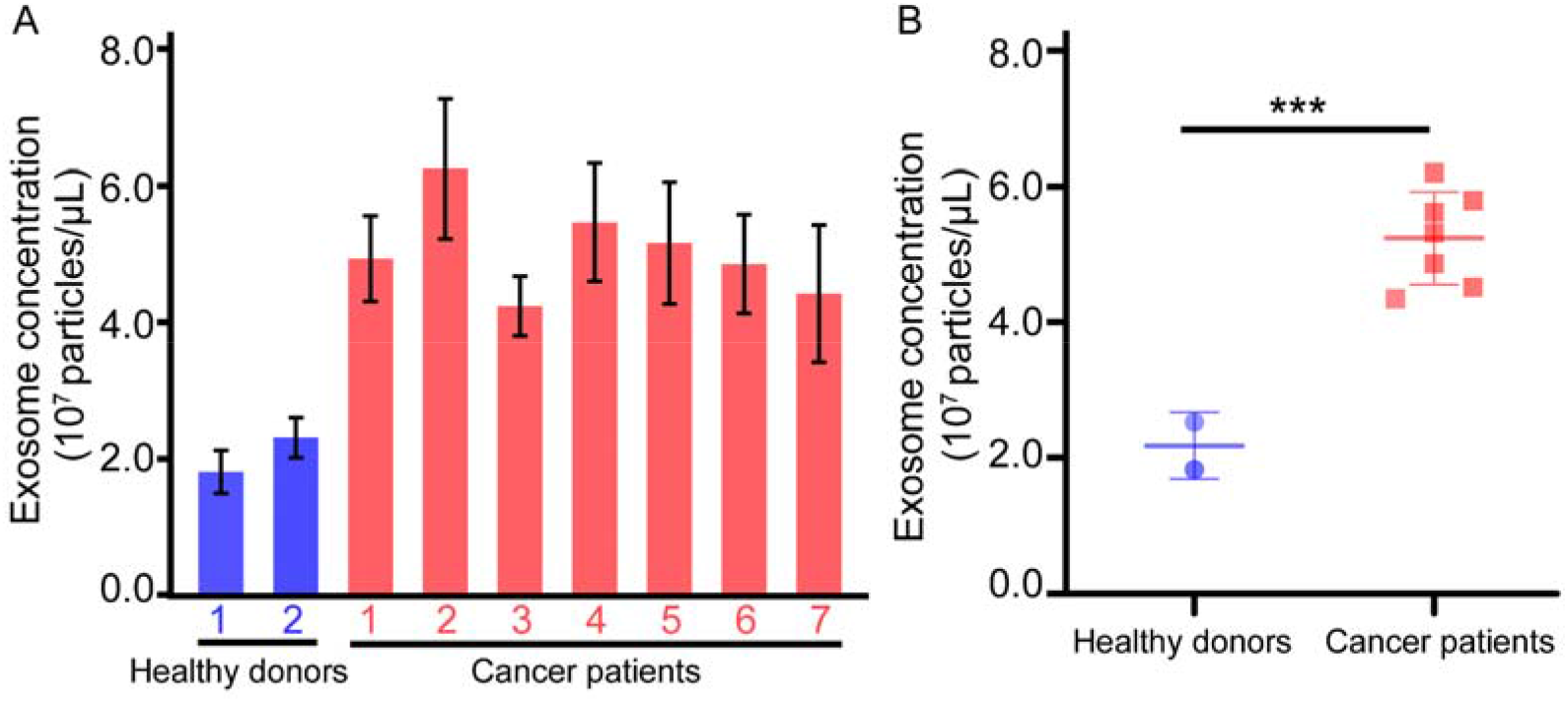
Determination of exosomes in serum samples from two healthy individuals and seven cancer patients. (A) Exosome concentrations of clinical samples analyzed by our method. (B) Scatter plot corresponding to panel A (*** p < 0.005).

## Conclusion

In conclusion, we proposed the use of DNA-nanostructures as a nanoweight to facilitate the enrichment of target exosomes by low-speed and short-term centrifugation, which not only simplifies the process but also maintains the integrity of exosomes. This method can specifically detect target MCF-7-derived exosomes in a dynamic range spanning 7 orders of magnitude, with an LOD as low as 1.8×10^2^ particles/μl, and be employed to analyze tumor-derived exosomes in human serum without requiring expensive reagents and equipment. We believe that our method represents a new example of DNA nanotechnology for applications in biomedical research and clinical diagnosis.

## Supporting information

Detailed experimental procedure, Western blotting analysis, characterization of HepG2-derived exosomes, comparison of the detection methods, and oligonucleotide sequences are supplied as Supporting Information.

## Acknowledgment

We thank the National Key Research and Development Program of China (2019YFA0900400), National Natural Science Foundation of China (21977046), Jiangsu Province Key Research and Development Program: Social Development Project (BE2021653), Program for Innovative Talents and Entrepreneur in Jiangsu, and Fundamental Research Funds for the Central Universities for financial support.

## TOC

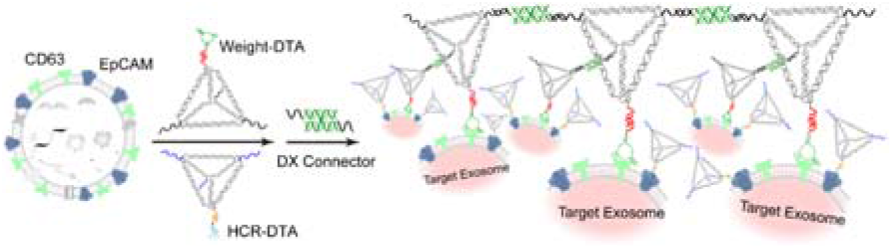

