## supplemental for "Fast and specific enrichment of cancer-related exosomes by DNA-nanoweight-assisted centrifugation"

**Experimental section**

**Reagent and material**

All DNA strands were purchased from Sangon Biotech. Co., Ltd. (Shanghai, China). The DNA sequences and modifications are listed in Table S2. Rabbit antihuman AFP, EpCAM and CD63 polyclonal antibodies were purchased from Proteintech Group Inc. (Chicago, IL). DyLight 800 4× PEG-conjugated secondary antibody was purchased from Cell Signaling Technology (Massachusetts, USA). Dulbecco’s modified Eagle medium (DMEM), fetal bovine serum (FBS), penicillin and streptomycin were purchased from Thermo Fisher Scientific Co., Ltd. (Waltham, MA). Albumin from bovine serum, sodium dodecyl sulfate (SDS), RIPA lysis buffer, trypsin-EDTA and other reagents were purchased from Sangon Biotech (Shanghai) Co., Ltd. Michigan Cancer Foundation-7 (MCF-7) and hepatocellular carcinoma cell line (HepG2) were purchased from Future Biotec (Nanjing, China). 30 kDa filters and 0.22 μm filters were purchased from Millipore Corp. (Bedford, MA). The solutions were prepared using deionized water purified by a Milli-Q water purification system (Millipore Corp., Bedford, MA).

The buffer solutions used in this work were as follows: (1) assembling buffer (1×TAE·Mg^2+^) containing 40 mM Tris, 40 mM acetic acid, 1 mM EDTA, and 12.5 mM Mg^2+^, pH 8.0. (2) Phosphate buffered saline (PBS) buffer containing 10 mM Na_2_HPO_4_, 10 mM NaH_2_PO_4_, 137 mM NaCl and 2.5 mM Mg^2+^, pH 7.4. (3) Western blot blocking buffer (TBST) containing 20 mM Tris·HCl, 300 mM NaCl, and 0.1% Tween 20, pH 7.4. (4) Washing buffer (PBST) containing 0.1% Tween-20 (v/v) in PBS buffer. (5) Transfer buffer containing 20 mM Tris, 100 mM glycine and 20% (v/v) methanol.

**Instrumentation.** The concentration of oligonucleotides was measured by a Nanodrop (Thermo Fisher Scientific Ltd., USA). Transmission electron microscopy (TEM) was performed using a JEOL JEM-2800 (JEOL Ltd., Japan). The concentration of exosomes was measured by nanoparticle tracking analysis (NTA) (XP Biomed Ltd., Shanghai). The size of exosomes was characterized by dynamic light scattering (DLS) (Malvern Zetasizer Nano ZS 90, UK). The fluorescence intensity was characterized by a microplate reader instrument (SpectraMax Id 3, Thermo Fisher).

**Assembly and purification of DNA nanostructure**. Weight-DTA was assembled from T1-A, T1-B, T1-C, T1-D and Apt-CD63, and HCR-DTA was assembled from T2-A, T2-B, T2-C, T2-D and Apt-EpCAM or Apt-AFP. Corresponding strands were mixed in equimolar ratio in 1×TAE·Mg^2+^ at a final concentration of 2 μM, heated to 95 °C for 2 min and cooled to 10 °C for 20 min. After the assembly of Weight-DTA and HCR-DTA, the 1×TAE·Mg^2+^ buffer was replaced with PBS by 30 kDa centrifugal filters, and excess aptamers were removed. The DX connector was mixed with M1 and M2 in equimolar ratios, heated to 95 °C for 2 min, and then slowly cooled to room temperature. To verify HCR amplification on 6% native PAGE, the initiator strand (0.1 μM), H1 and H2 were mixed at a ratio of 1:3:15 for 2 hours.

**Collection of crude exosome**. Crude exosomes were collected from HepG2 and MCF-7 cells using conventional centrifugation. At a cell density of 80%-90%, the cells were cultured in DMEM without 10% FBS for 24 hours. The cell culture supernatant was collected by a series of centrifugations at 4 °C with a speed of 300×g for 10 min, 2,000×g for 10 min, and 10,000×g for 30 min in sequence to discard cellular debris. Capsule fragments were then removed by a 0.22 μm filter.

**Exosome enrichment.** For Weight-DTA-assisted centrifugation, 2 μM 10 μL Weight -DTA was added to 180 μL crude exosome solution (approximately 1.7×10^10^ particles per milliliter as measured by NTA, the rate of Weight -DTA and exosome was about 3900:1) and incubated at room temperature for 120 min. Then, 6 μM 10 μL DX connector was added and incubated for 1 hour. After centrifugation at 20,000×g for 30 minutes, target exosome pellets were collected. For conventional ultracentrifugation, crude exosome solution was centrifuged at 140,000×g for 90 min at 4 °C. After the resuspension of pellets in PBS, another centrifugation followed at 140,000× g for 90 min at 4 °C. Finally, the pellets were resuspended in 300 μL PBS.

**HCR amplification.** Ten microliters of 1 μM purified HCR-DTA was added to 90 μl of enriched exosomes and incubated at room temperature for 30 min. The supernatant was removed by centrifugation at 15,000×g for 30 min, and the pellets were resuspended in 90 μL of PBS. After the addition of 10 μL 3 μM H1 and 10 μL 15 μM H2 for 2 hours at room temperature, the emission spectrum was measured by a fluorescence microplate reader.

**Analysis by Western Blotting.** The target proteins of the exosomes were lysed by a total protein extraction kit and separated by SDS–PAGE. The protein bands were transferred to fluoride membranes (PVDF) by Western Semidry Transfer and blocked with 5% (w/v) BSA in TBST buffer for 2 hours at room temperature. The corresponding primary antibodies were added and incubated overnight at 4 °C. After washing three times with TBST buffer, the membrane was incubated with DyLight 800-modified secondary antibody for 1 hour at room temperature. The band was obtained by ODYSSEY CLX System.

**Exosome Detection in Human Serum Samples.** To investigate the clinical application of our methods for the enrichment and detection of exosomes, human serum samples of breast cancer patients and healthy individuals were obtained from the Jiangsu Province Hospital. Written informed consent was obtained from all study participants prior to enrollment, and the study was approved by the ethics committees from each institution involved. Prior to analysis, the samples were centrifuged at 3,000 g for 10 min and filtered by a 0.22 μm filter. Each serum sample was diluted to 200 μL for exosome detection. The detection procedures for exosome in human serum was the same as the procedure for exosome detection described above.

**Results and Discussion**

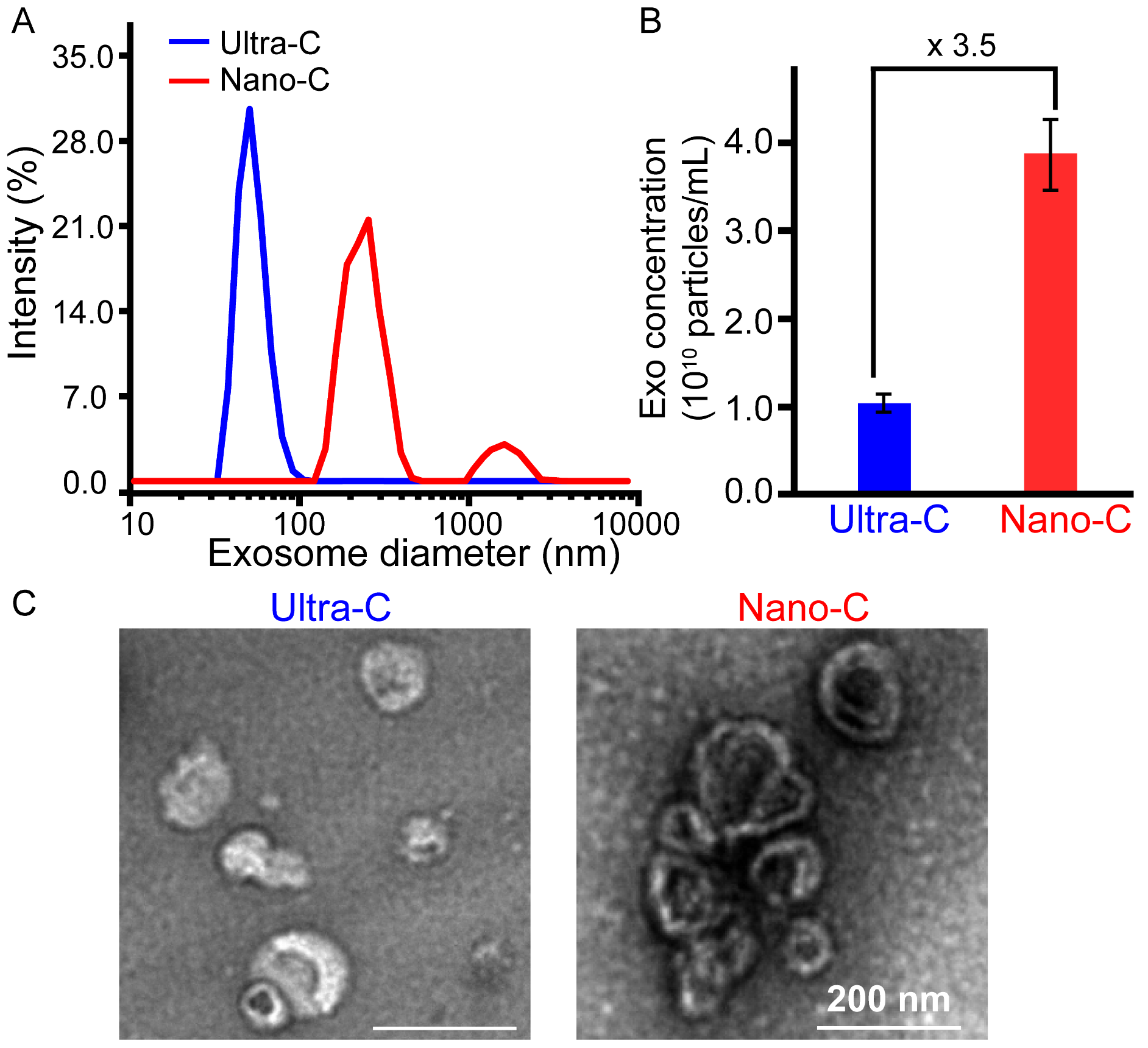

**Figure S1**. Enrichment of HepG2 derived exosomes by Nanoweight-assisted centrifugation (Nano-C). (A) DLS analysis, (B) total protein amount and (C) TEM images of HepG2 derived exosomes isolated by Nano-C or conventional ultracentrifugation (Ultra-C) Data are represented as the means ± SD (n = 4).

Table S1. Comparison of our method and recently reported methods of exosome detection

| Method | LOD (particles/μL) | Enrichment time | Quantification time |
| --- | --- | --- | --- |
| Ref 1 | 1.4×10^5^ | 1 h | 1 h |
| Ref 2 | 3.96×10^2^ | > 3 h | 30 min |
| Ref 3 | 1.8×10^2^ | > 3 h | 1 h |
| Ref 4 | 8.2×10^3^ | > 3 h | 1 h |
| Ref 5 | 3.0×10^4^ | > 3 h | 30 min |
| Ref 6 | 2.08×10^5^ | > 3 h | 1 h |
| Ref 7 | 3.0×10^4^ | NA ^a)^ | 1 h |
| Ref 8 | 3.4×10^6^ | NA | 15 min |
| Ref 9 | 6.56×10^4^ | NA | 1 h |
| This work | 1.8×10^2^ | 40 min | 1 h |

NA ^a)^ refers to not available

Table S2. Sequences of oligonucleotides used in this work

| Name | Sequence (5’- 3’) |
| --- | --- |
| T1-A | ACATTCCTAAGTCTGAAACATTACAGCTTGCTACACGAGAAGAGCCGCCATAGTAAGCAACCTGCCTGTTAGC |
| T1-B | TATCACCAGGCAGTTGACAGTGTAGCAAGCTGTAATAGATGCGAGGGTCCAATACTCTCTTCACCGTAATCTT |
| T1-C | TCAACTGCCTGGTGATAAAACGACACTACGTGGGAATCTACTATGGCGGCTCTTCTCTCTTCACCGTAATCTT |
| T1-D | TTCAGACTTAGGAATGTGCTTCCCACGTAGTGTCGTTTGTATTGGACCCTCGCATTCTCTTCACCGTAATCTT |
| M1 | GACCCTAAGCATACATGATTACGGTGAAGAGA |
| M2 | ATGTATGCTTTAGGGTC |
| Apt-CD63 | CACCCCACCTCGCTCCCGTGACACTAATGCTATTTTGCTAACAGGCAGGTTGCT |
| T2-A | AGTCTAGGATTCGGCGTGGGTTAAACATTCCTAAGTCTGAAACATTACAGCTTGCTACACGAGAAGAGCCGCCATAGTA |
| T2-B | AGTCTAGGATTCGGCGTGGGTTAATATCACCAGGCAGTTGACAGTGTAGCAAGCTGTAATAGATGCGAGGGTCCAATAC |
| T2-C | AGTCTAGGATTCGGCGTGGGTTAATCAACTGCCTGGTGATAAAACGACACTACGTGGGAATCTACTATGGCGGCTCTTC |
| T2-D | GCCGTCGTGCCTTATTTCTGTTCAGACTTAGGAATGTGCTTCCCACGTAGTGTCGTTTGTATTGGACCCTCGCAT |
| Apt-EpCAM | CAGAAATAAGGCACGACGGCCACTACAGAGGTTGCGTCTGTCCCACGTTGTCATGGGGGGTTGGCC |
| Apt-AFP | CAGAAATAAGGCACGACGGCGTGACGCTCCTAACGCTGACTCAGGTGCAGTTCTCGACTCGGTCTTGATGTGGGTCCTGTCCGTCCGAACCAATC |
| H1 | TTAACCCACGCCGAAT/BHQ/CCTAGACTCAAAGTAGTCTAGGA/Fam/TTCGGCGTG |
| H2 | AGTCTAGGATTCGGCGTGGGTTAACACGCCGAATCCTAGACTACTTTG |
